# Machine learning on large scale perturbation screens for SARS-CoV-2 host factors identifies β-catenin/CBP inhibitor PRI-724 as a potent antiviral

**DOI:** 10.1101/2023.02.23.529833

**Authors:** Maximilian A. Kelch, Antonella Vera-Guapi, Thomas Beder, Marcus Oswald, Alicia Hiemisch, Nina Beil, Piotr Wajda, Sandra Ciesek, Holger Erfle, Tuna Toptan, Rainer König

## Abstract

Expanding antiviral treatment options against SARS-CoV-2 remains crucial as the virus evolves rapidly and drug resistant strains have emerged. Broad spectrum host-directed antivirals (HDA) are promising therapeutic options, however the robust identification of relevant host factors by CRISPR/Cas9 or RNA interference screens remains challenging due to low consistency in the resulting hits.

To address this issue, we employed machine learning based on experimental data from knockout screens and a drug screen. As gold standard, we assembled perturbed genes reducing virus replication or protecting the host cells. The machines based their predictions on features describing cellular localization, protein domains, annotated gene sets from Gene Ontology, gene and protein sequences, and experimental data from proteomics, phospho-proteomics, protein interaction and transcriptomic profiles of SARS-CoV-2 infected cells.

The models reached a remarkable performance with a balanced accuracy of 0.82 (knockout based classifier) and 0.71 (drugs screen based classifier), suggesting patterns of intrinsic data consistency. The predicted host dependency factors were enriched in sets of genes particularly coding for development, morphogenesis, and neural related processes. Focusing on development and morphogenesis-associated gene sets, we found β-catenin to be central and selected PRI-724, a canonical β-catenin/CBP disruptor, as a potential HDA. PRI-724 limited infection with SARS-CoV-2 variants, SARS-CoV-1, MERS-CoV and IAV in different cell line models. We detected a concentration-dependent reduction in CPE development, viral RNA replication, and infectious virus production in SARS-CoV-2 and SARS-CoV-1-infected cells. Independent of virus infection, PRI-724 treatment caused cell cycle deregulation which substantiates its potential as a broad spectrum antiviral. Our proposed machine learning concept may support focusing and accelerating the discovery of host dependency factors and the design of antiviral therapies.

**Author’s summary:** Drug resistance to pathogens is a well-known phenomenon which was also observed for SARS-CoV-2. Given the gradually increasing evolutionary pressure on the virus by herd immunity, we attempted to enlarge the available antiviral repertoire by focusing on host proteins that are usurped by viruses. The identification of such proteins was followed within several high throughput screens in which genes are knocked out individually. But, so far, these efforts led to very different results. Machine learning helps to identify common patterns and normalizes independent studies to their individual designs. With such an approach, we identified genes that are indispensable during embryonic development, i.e., when cells are programmed for their specific destiny. Shortlisting the hits revealed β-catenin, a central player during development, and PRI-724, which inhibits the interaction of β-catenin with cAMP responsive element binding (CREB) binding protein (CBP). In our work, we confirmed that the disruption of this interaction impedes virus replication and production. In A549-AT cells treated with PRI-724, we observed cell cycle deregulation which might contribute to the inhibition of virus infection, however the exact underlying mechanisms needs further investigation.

## Introduction

Genetic plasticity of SARS-CoV-2 and herd immunity-derived evolutionary pressure have led to the emergence of variants of concern (VOC) (1, 2). Some of the variants and recombinant lineages have evaded vaccine-elicited antibodies and monoclonal antibody-based treatments (3–6). Although antiviral drugs such as Remdesivir, Molnupiravir, or Nirmatrelvir/Ritonavir (Paxlovid) have significantly improved disease outcome in patients (7–10), these treatments might select for more resistant variants over time (11, 12). Consequently, our current virus-directed countermeasures have limitations and there is an urgent need for developing host-directed antivirals (HDA) that can target host-dependency factors (HDF) required for the life cycle of the virus. HDA might provide variant-independent treatment opportunities to reduce disease severity and complement virus-directed antiviral therapies (13–15). In addition, drug repurposing can offer an expedited timeline to bring HDA therapies into clinical settings in a cost-effective and timely manner. Still, a common challenge is the robust identification of relevant host factors among varying *in vitro* conditions, although recent advances in functional genomics have simplified the use of high-throughput technology platforms based on CRISPR/Cas9 and siRNAs (16–19). However, when comparing the results of these experimental screens for SARS-CoV-2 host dependency factors, there is only a marginal overlap of hits. This may be due to different viral strains, host cells and/or different multiplicity of infection (MOI) or further different experimental settings such as different knockout library constructs. Interestingly, when grouping the identified HDF into gene sets of common function or cellular processes, the consistency increases suggesting that these lists of HDF in their specific contexts contain consistent patterns on a more complex level.

Machine learning methods can identify such common patterns in experimental data, as e. g. the identification of common regulators explaining large scale transcription profiles (20, 21) and allow for integration of broad variety of omics and genomics data. In this study, we followed a machine learning approach to identify HDF of SARS-CoV-2. From the knockout screens (16–19), we assembled a gold standard of genes required for virus replication. With this gold standard, we trained and evaluated the classifier. The classifier based its predictions on features of the genes describing their encoded proteins’ cellular localization, protein domains, annotated cellular processes, and gene/protein sequences. Furthermore, we embedded experimental data comprising proteomic, PPI, and transcriptomic profiles of SARS-CoV-2 infected cells. In addition, another classifier was trained based on a gold standard derived from a drug screen (22), in which more than 6,000 drugs were observed if they provide a protective effect on SARS-CoV-2 infected cells. Both classifiers delivered a list of predicted HDF. For a few, well selected HDF, we interrogated drug databases, and identified β-catenin and its small molecule inhibitor PRI-724, particularly antagonizing the interaction between β-catenin and CBP.

The β-catenin/CBP interaction displays a pivotal part of a switch-like signaling network within the Wnt/β-catenin pathway that controls the well-balanced interplay between cell proliferation and differentiation (23, 24). Activation of Wnt signaling leads to a sequester of the β-catenin destruction complex enabling cytosolic accumulation of β-catenin and, in turn, its nuclear translocation where it interacts with CBP or p300 to drive either proliferation or differentiation, respectively (25). In this study, we identified β-catenin as a SARS-CoV-2 host factor using machine-learning and sought to analyze the effectiveness of one of its inhibitors, PRI-724, against pandemic RNA viruses including SARS-CoV-2 in different cell line models.

## Results

### Host dependency factors identified by genome scale screens showed marginal overlap on the gene level, but improved consistency for common cellular processes and functions

First, the overlap of the results from published screening studies were investigated, in the following denoted as Daniloski *et al*., Zhu *et al*., Wei *et al*., and Wang *et al*. (16–19). For a balanced comparison, we regarded the top 500 genes with the highest scores of every screen (**Fig 1A**). Only ACE2 was common among all candidates. The highest overlap (n = 37 genes) was observed between the screens of Daniloski *et al*., and Zhu *et al*.. Besides ACE2, only 11 genes were common among Wang *et al*., and Wei *et al*.. All these overlaps were not significantly enriched (using Fisher’s Exact Test). Similarly, we found very low correlations while analyzing the ranking of all genes of the screens (independent of the cutoff) (**Fig 1B**). Next, we investigated how grouping of genes into gene sets of common cellular processes and gene functions affect the results. For this, gene set enrichment tests were performed with the same lists of the 500 highest scoring genes (**Fig 1C**). Interestingly, the overlaps were considerably higher but still not significant. Daniloski *et al*. and Zhu *et al*. shared 44/51 gene sets, Daniloski *et al*., Wang *et al*., and Zhu *et al*. shared 5/51 gene sets and 2/51 were identical among Daniloski *et al*., Wei *et al*., and Zhu *et al*.. The overlapping gene sets included vesicle-mediated transport to the plasma membrane, endosomal and lysosomal transport, phagosome maturation, transforming growth factor-beta production, post-Golgi vesicle-mediated transport, transition metal ion transport, negative regulation of cell growth, and phosphatidylinositol biosynthetic process (**Suppl. Tab S1**). In summary, we observed limited overlap among the investigated screens. The overlap was higher when regarding gene sets of common cellular processes or molecular function. This supported our concept to set up a machine learning procedure to identify more common patterns among these different datasets.

**Fig 1:**
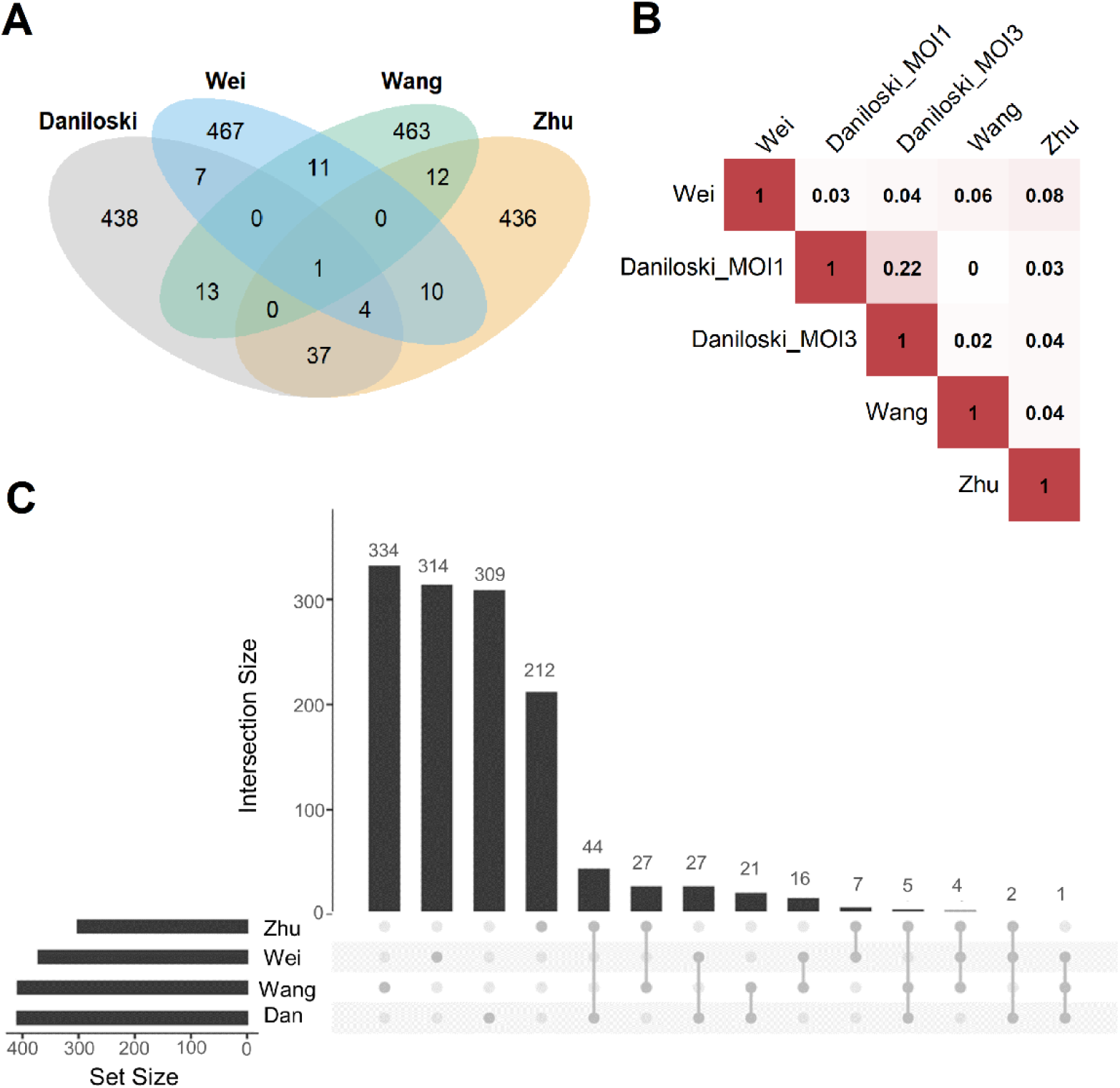
Overlap analysis of publicly available host factor identification screens. **(A)** Intersections of the experimentally obtained HDF (500 genes with the highest scores in the screens) among the different knockout screening studies. Only ACE2 (n=1, center of the figure) was identified as an HDF in all screens. **(B)** Pairwise correlations between the ranking of all commonly screened genes of the four studies (Daniloski *et al*., Zhu *et al*., Wang *et al*., Wei *et al*.). Red represents a positive correlation, white no correlation, Daniloski_MOI1: data of the Daniloski screen with MOI=0.01, Daniloski_MOI3: data of the Daniloski screen with MOI=0.3. **(C)** This diagram (upset plot) shows the overlaps of gene sets being enriched of HDF (again considering the 500 genes with the highest scores in the screens), Dan: Daniloski.

### Classifiers based on data from the knockout screens and a drug based screen performed well in predicting HDF from the gold standards

We assembled a list of 500 genes commonly high ranking among the investigated knockout screens (top 500 genes of the rank products, see Methods). This list served as the input for the machine learning classifier. Similarly, we selected negative controls from the lowest ranking genes. Furthermore, we set up a descriptor for gene predictions by compiling a comprehensive list of more than 60,000 features for each gene describing its nucleic acid and protein sequences, potential associated cellular processes, its associated compartments, functional domains, molecular functions, its conservation, and its network topology within a protein interaction network. Here, also other omics data of SARS-CoV-2 infected cells, like gene transcription profiles, proteomics and interactions between viral and host proteins were integrated. Performing a cross-validation, in which training and validation data is strictly separated, we observed good performance results with an area under the curve (AUC) of a Receiver Operator Characteristics of 0.82 (1σ = 0.03) (**Fig 2A**). In addition, the same procedure was performed and a classifier learned based on data of a drug screen (in the following drug screen based classifier) (22). Here, we obtained slightly worse performance compared to the knockout-based classifier receiving an AUC of the Receiver Operator Characteristics of 0.71 (1σ=0.11). Additional performance parameters are supplemented (**Suppl. Tab S2**).

**Fig 2:**
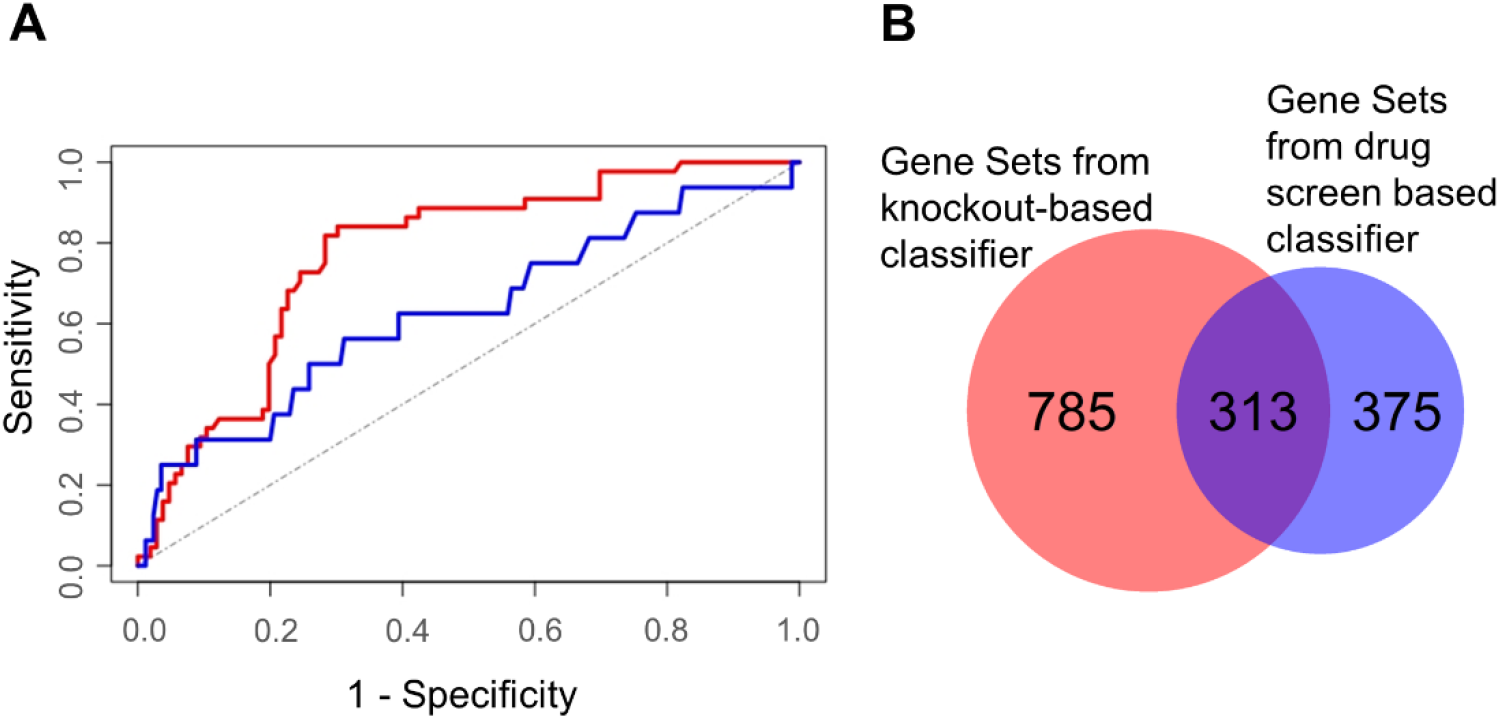
Characterization of machine learning performance and overlap of the gene sets derived from the two classifier approaches. **(A)** Machine learning performance results depicted by the Receiver Operator Characteristics of the validation sets. The area under the curve was 0.82 for the knockout screens based classifier (red) and 0.71 for the drug screen based classifier (blue). The (theoretical) result from random guessing is indicated by the dotted line, **(B)** Gene sets being enriched in HDF predicted by the knockout screening based classifier (left) and the drug screening based classifier (right).

### Predicted HDF are prominently enriched in gene sets related to morphogenesis and development

We selected 2,182 and 1,989 top-scoring genes identified by knockout- and drug screen-based classifiers, respectively. To identify the most involved cellular mechanisms in the life cycle of the virus, we performed gene set enrichment analysis for each list illuminating 1,098 and 688 gene sets of the respective classifiers (**Suppl. Tab S3 and S4**). Interestingly, we observed a high number of common gene sets (n = 313) showing consistency among both classifiers. Regarding these common gene sets, we further found a high fraction of these gene sets (69 out of 313 gene sets) to be related to morphogenesis and development, followed by 60 gene sets being related to neural processes (**Suppl. Tab S5**). Besides these gene sets, further gene sets were related to signaling, gene regulation, and immune and stress response (**Suppl. Tab S5**).

During development, a large variety of distinct different cellular identities need to be established and maintained in the embryo. Particularly, during developmental lineage reprogramming, a somatic cell can be reprogrammed into a distinct cell type by forced expression of lineage-determining factors (76). Similarly, coronaviruses reprogram their host cell for their specific needs, including their replication (26–30). We followed this interesting parallelism and selected a gene list from the gene sets of development and morphogenesis for further prioritization. Aiming to interfere “reprogramming” with a high impact, we prioritized genes coding for proteins with high connectivity in a protein-protein interaction (PPI) network to enhance the impact of a treatment by small molecule inhibitors. This led to a short list of genes with the highest connectivity (number of nearest neighbors) in this PPI network (**Tab 1** lists the top ten). Out of these, β-catenin (CTNNB1) was selected for further analysis due to its high gene expression in SARS-CoV-2 infected cells. Interrogating publicly available compound and drug databases led to the selection of PRI-724 for experimental follow up described in the next sections.

**Tab 1.**
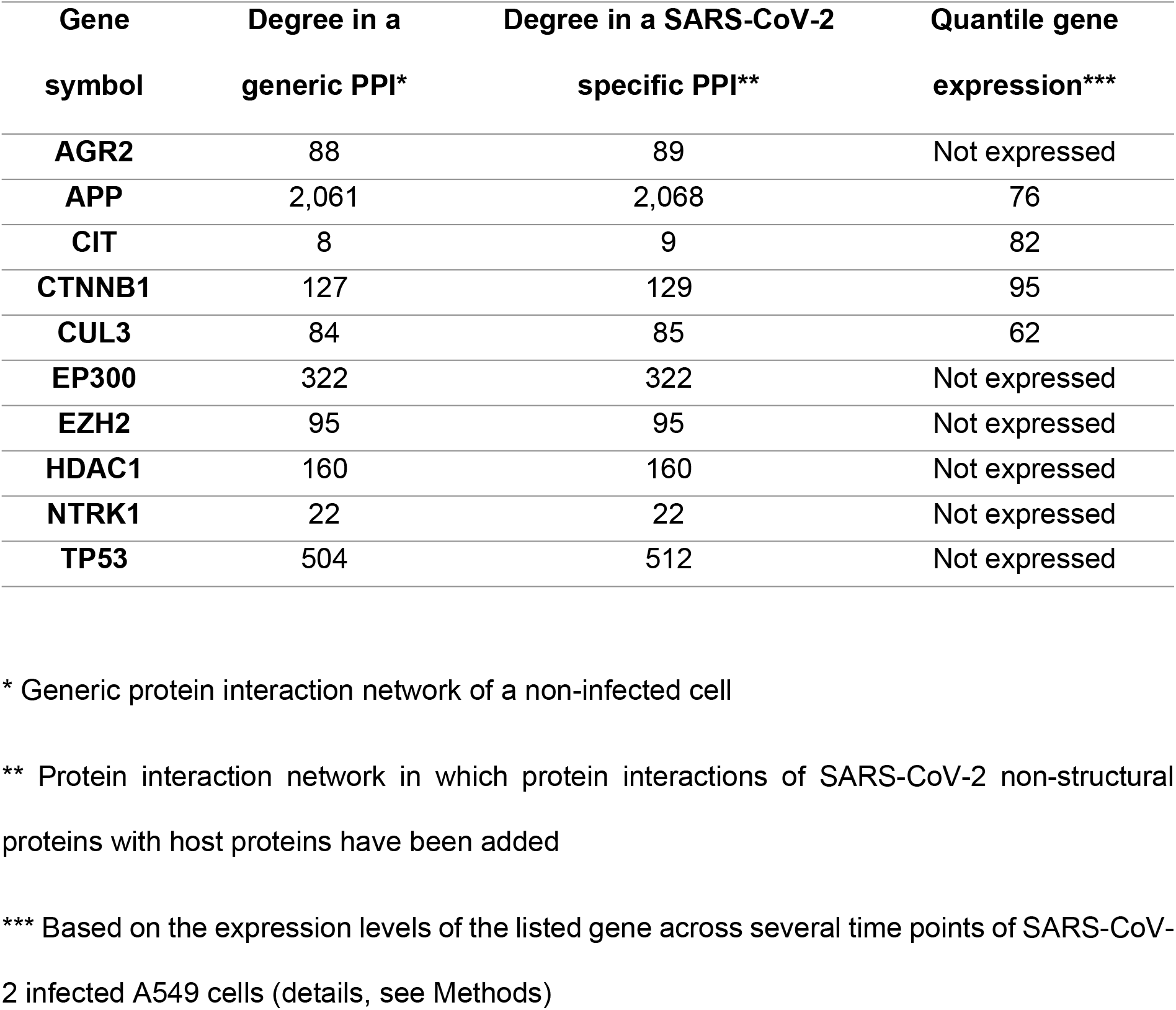
Short list of selected genes of morphogenesis and development with high connectivity.

| Gene<br>symbol | Degree in a<br>generic PPI* | Degree in a SARS-CoV-2<br>specific PPI** | Quantile gene<br>expression*** |
| --- | --- | --- | --- |
| AGR2 | 88 | 89 | Not expressed |
| APP | 2,061 | 2,068 | 76 |
| CIT | 8 | 9 | 82 |
| CTNNB1 | 127 | 129 | 95 |
| CUL3 | 84 | 85 | 62 |
| EP300 | 322 | 322 | Not expressed |
| EZH2 | 95 | 95 | Not expressed |
| HDAC1 | 160 | 160 | Not expressed |
| NTRK1 | 22 | 22 | Not expressed |
| TP53 | 504 | 512 | Not expressed |

\* Generic protein interaction network of a non-infected cell
\*\* Protein interaction network in which protein interactions of SARS-CoV-2 non-structural proteins with host proteins have been added
\*\*\* Based on the expression levels of the listed gene across several time points of SARS-CoV-2 infected A549 cells (details, see Methods)

### PRI-724 inhibits SARS-CoV-2 variants, SARS-CoV-1, and Influenza A virus in A549-AT cells

In order to validate *in silico* findings regarding PRI-724 in the context of an infection, various SARS-CoV-2 isolates (Ancestral variants: B (FFM5), B.1 (FFM7, D614G); variants of interest (VOI): P.2 (Zeta), B.1.429 (Epsilon), B.1.617.1 (Kappa); deescalated variants of concern (VOC): B.1.1.7 (Alpha), B.1.351 (Beta), B.1.617.2 (Delta); VOC: B.1.1.529 BA.1, BA.2, and BA.5 (Omicron)), and SARS-CoV-1 were tested. A549-AT cells were infected at an MOI of 0.1 and treated with PRI-724 or Remdesivir for 48 h. CPE-related confluency changes were measured using automated label-free brightfield microscopy (**Fig 3A**). For all viruses tested, a mean IC_50_ of 1.491 μM (95%CI 1.087-1.896) and IC_50_ of 0.2916 μM (95%CI 0.1948-0.3885) was observed for PRI-724 and Remdesivir, respectively. No significant differences were detected among SARS-CoV-2 variants, even though the B.1 and B.1.429 variants showed ~1.5-fold and ~2-fold higher IC_50_, respectively, and BA.1, BA.2, and BA.5 variants had ~2-fold lower IC_50_ (**Fig 3A, Suppl. Tab S6**). The overall response pattern among different variants remained similar for Remdesivir. Treatment with the PRI-724 active metabolite C-82 and the analog ICG-001 likewise inhibited SARS-CoV-2 (B.1.617.2 isolate) in A549-AT with IC_50_ values of 1.005 μM and 14.85 μM, respectively (**Suppl. Fig S1A**). In addition to that, dose-responses to PRI-724 for SARS-CoV-2 variants B.1, B.1.1.7, B.1.351, P.2, and B.1.617.2 were obtained in Calu-3 cells, with overall increased IC_50_ of 8.448 μM (95%CI 7.431-9.465) for PRI-724 and mean IC_50_ of 0.4962 (95%CI 0.2686-0.7237) for Remdesivir (**Suppl. Fig S1B, Suppl. Tab S7**). In Calu-3, PRI-724 further demonstrated inhibition of MERS-CoV (IC_50_ = 22.4 μM) and showed significantly decreased SARS-CoV-1 nucleocapsid (N) expression upon treatment with 10 μM, 30 μM, and 100 μM PRI-724 (**Suppl. Fig S1C**). In an attempt to validate our results and to test whether PRI-724 can be used against a broader spectrum of RNA viruses, we infected A549-AT cells with Influenza A virus (IAV), SARS-CoV-2 (B.1.617.2 strain), or SARS-CoV-1, and performed immunofluorescence staining for IAV nucleoprotein (NP) and the coronavirus N (**Fig 3B**). Significant concentration-dependent reduction of NP expression was evident for treatment with 10 μM, 3 μM, and 1 μM PRI-724. Treatment with 10 μM and 3 μM PRI-724 led to ~10-fold and ~4-fold reduction of H5N1 NP^+^ cells, whereas for these treatments no cells stained positive for SARS-CoV-1 N. Treatment with 3 μM reduced SARS-CoV-2 N^+^ cells ~15-fold and staining was negative with 10 μM PRI-724.

**Fig 3:**
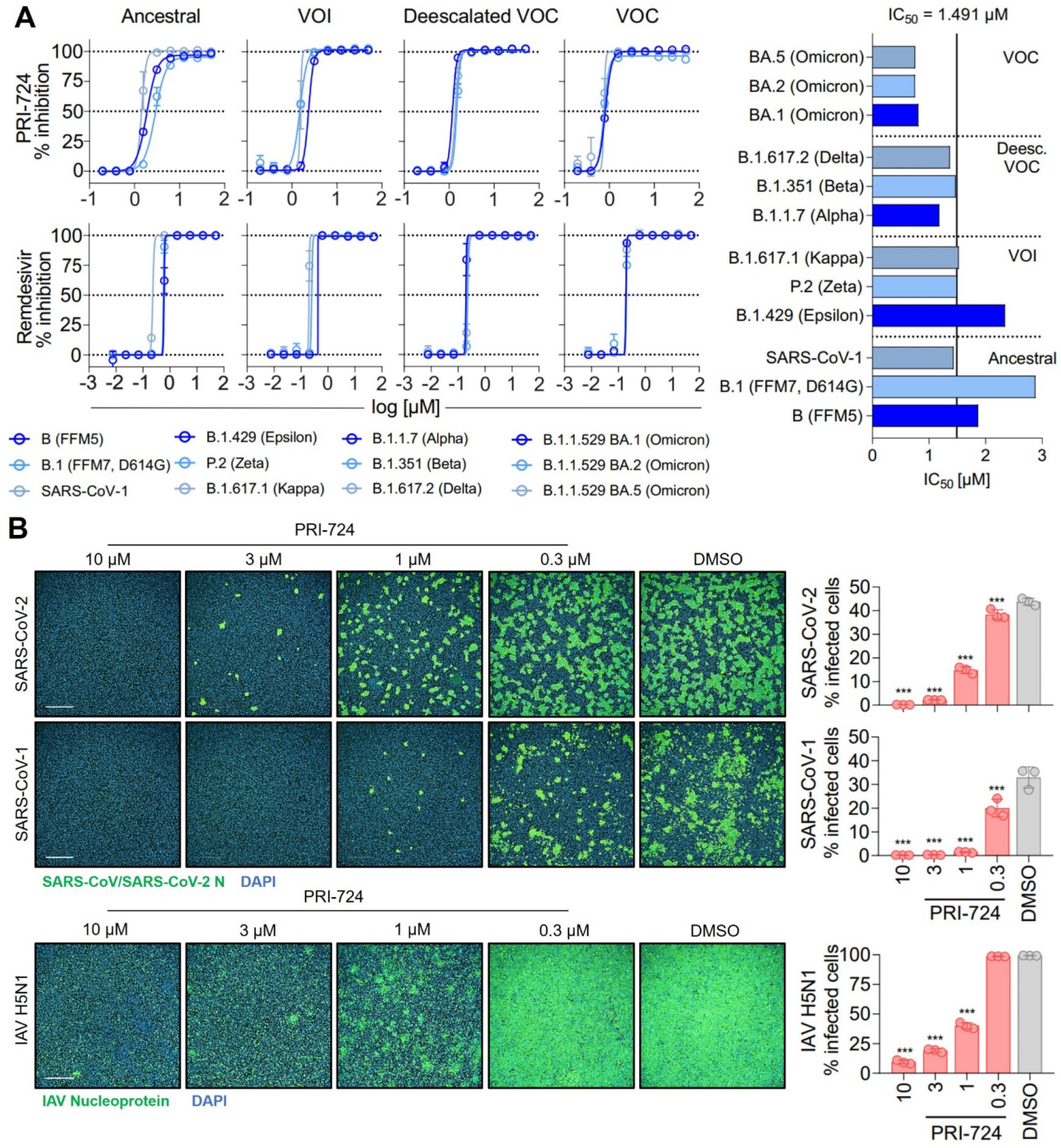
Inhibition of SARS-CoV-2, SARS-CoV-1 and IAV H5N1 by PRI-724. **(A)** Dose-response curves of diverse SARS-CoV-2 mutational variants and SARS-CoV-1 to PRI-724 and Remdesivir. Cell confluence was analyzed 48 hpi using a Spark^®^ Cyto Imaging System. Dose-response curves were fit to normalized % inhibition. Data represent mean ± SD of three biological replicates. The experiment was repeated twice with similar results. IC_50_ values were calculated by applying robust non-linear regression. Mann-Whitney-U test confirmed no significant changes between any dose-responses analyzed. **(B)** Staining of SARS-CoV-2 (B.1.617.2 strain) and SARS-CoV-1 N protein, as well as IAV H5N1 NP 18 hpi. Data points represent mean ± SD of three biological replicates. The experiment was repeated twice with similar results. Infected cells were determined by Alexa 488/DAPI co-localization using an Operetta^®^ High-Throughput Imaging system. Scale bars = 500 μm.

### PRI-724 inhibits syncytium formation and CPE onset in A549-AT cells

After confirming PRI-724-mediated inhibition of CPE in A549-AT cells at 48 hpi, we examined the timeline of CPE onset and development. Label-free live-cell brightfield imaging was implemented to address viral growth kinetics upon treatment with PRI-724. Cells were treated with 3 μM (high-dose), 1 μM (low-dose) PRI-724, and the highest corresponding amount of DMSO and were subsequently infected with SARS-CoV-2 (B.1.617.2 isolate) or SARS-CoV-1 (MOI=0.1). Imaging, confluence measurements and surface roughness measurements were performed over the course of 48 h. A549-AT confluence as well as surface roughness were mostly comparable among all treatment conditions until 24 hpi for both viruses. After 24 hpi, virus-induced cell lysis gradually reduced the confluence and increased surface roughness in DMSO-treated cells. SARS-CoV-2 and SARS-CoV-1 infected cells treated with low-dose PRI-724 started to lyse at 29 hpi and 39 hpi, respectively, while high-dose PRI-724 treated cells showed no substantial loss in confluence over the course of the experiment (**Fig 4A, Videos S1-S6**). These findings were confirmed by brightfield microscopy at indicated time points (**Fig 4B**). Here, cell-cell-fusion, i. e. syncytium formation, caused by both viruses, was initially detectable at 16 hpi and 24 hpi, respectively, in DMSO-treated cells and was not present in high-dose PRI-724 treated cells. In low-dose PRI-724 treated cells, SARS-CoV-2 syncytia and CPE were detectable, but much smaller compared to the control (**Fig 4B, row 2-3**). Although, we found likewise results for SARS-CoV-1, cell lysis initiated at 32 hpi compared to the control (**Fig 4B, row 5-6**).

**Fig 4:**
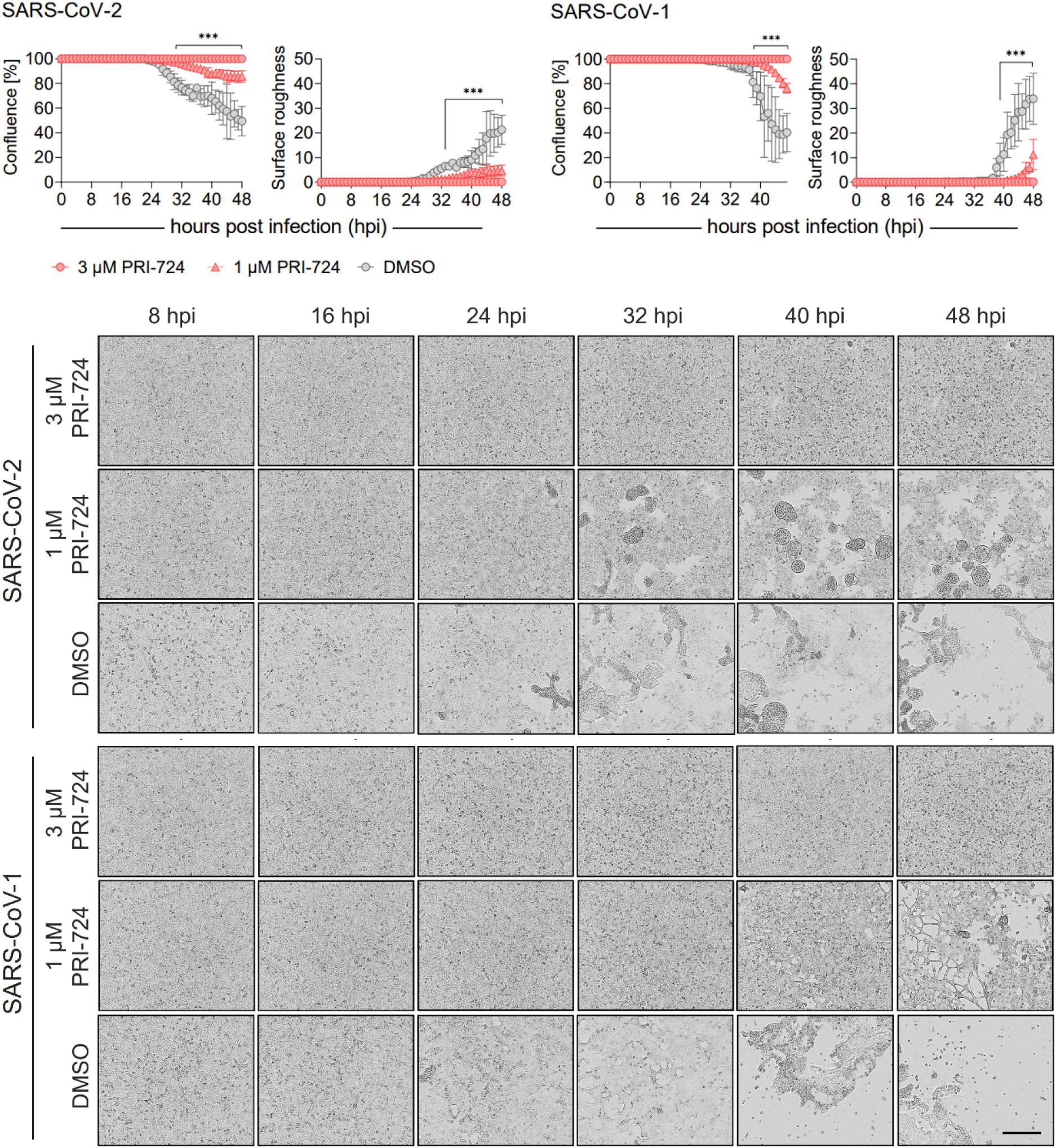
Growth kinetics of PRI-724 treated A549-AT cells. A549-AT cells treated with high (3 μM) and low (1 μM) dose of PRI-724 were subsequently infected with SARS-CoV-2 (B.1.617.2 isolate) and SARS-CoV-1 (FFM1 isolate) (MOI=0.1). Cells were incubated at 37°C and 5% CO_2_ for 48 h in a Spark^®^ Cyto Imaging system. Graphical presentation of % confluence and surface roughness for both SARS-CoV-2 and SARS-CoV-1 are shown for measurements hourly performed. Respective brightfield microscopy images represent critical timepoints of CPE formation over the course of infection during treatment with PRI-724 and without. Scale bar: 200 μm. See also **Videos S1-S6**.

Every condition comprises three biological replicates and graphs represent means ± SD; significant differences (\*\*\**p*<0.001) are indicated by asterisks obtained by performing multiple t tests. The experiment was performed twice yielding similar results.

### PRI-724 reduces virus RNA replication and infectious virus progeny

Since PRI-724 inhibited CPE development, we questioned whether it impaired virus replication and/or progeny virus production. For the replication analysis, the expression of intracellular subgenomic nucleocapsid (sg-N) RNA was monitored over the course of 24 h and 48 h in A549-AT and Calu-3 cells, respectively (**Fig 5A**). In A549-AT cells, intracellular SARS-CoV-2 (B.1.617.2 isolate) sg-N RNA was significantly reduced at 6, 10, 12, and 24 hpi when treated with 10 μM, 3 μM, and 1 μM PRI-724 just prior to infection. Most remarkable changes were detected 6 hpi and 24 hpi where vRNA was reduced ~370-fold/~96-fold, ~38-fold/~13-fold, and ~20-fold/~7-fold (**Suppl. Tab S8**) compared to the control when treated with 10 μM, 3 μM, and 1 μM PRI-724, respectively. Similarly, in Calu-3, SARS-CoV-2 vRNA was significantly reduced at 8, 36, and 48 hpi, with most noticeable changes at 36 hpi, namely ~105-fold, ~69-fold, and ~10-fold (**Suppl. Tab S8**) after treatment with 20 μM, 15 μM, and 10 μM PRI-724, respectively. For SARS-CoV-1 in A549-AT, PRI-724 treatment reduced vRNA at 8, 10, 12, and 24 hpi. In comparison to SARS-CoV-2, the impact on SARS-CoV-1 vRNA levels was more pronounced. 24 hpi SARS-CoV-1 vRNA was reduced ~1,288-fold, ~157-fold, and ~22-fold (**Suppl. Tab S8**) upon treatments with 10 μM, 3 μM, and 1 μM PRI-724, respectively. In A549-AT and Calu-3, Remdesivir treatment appeared more effective in terms of vRNA reduction compared to PRI-724, but timepoints of significant reductions were comparable between both treatments. After 24 h in A549-AT, Remdesivir reduced SARS-CoV-2 and SARS-CoV-1 vRNA 270-fold and ~3,142-fold, respectively (**Suppl. Tab S8**). In Calu-3, SARS-CoV-2 RNA was reduced ~249-fold (**Suppl. Tab S8**) at 24 hpi upon treatment compared to the DMSO control. (**Fig 5A**). In general, reduced sg-N copies correlated with lower total N RNA in A549-AT and Calu-3 cells. (**Suppl. Fig S2**).

**Fig 5:**
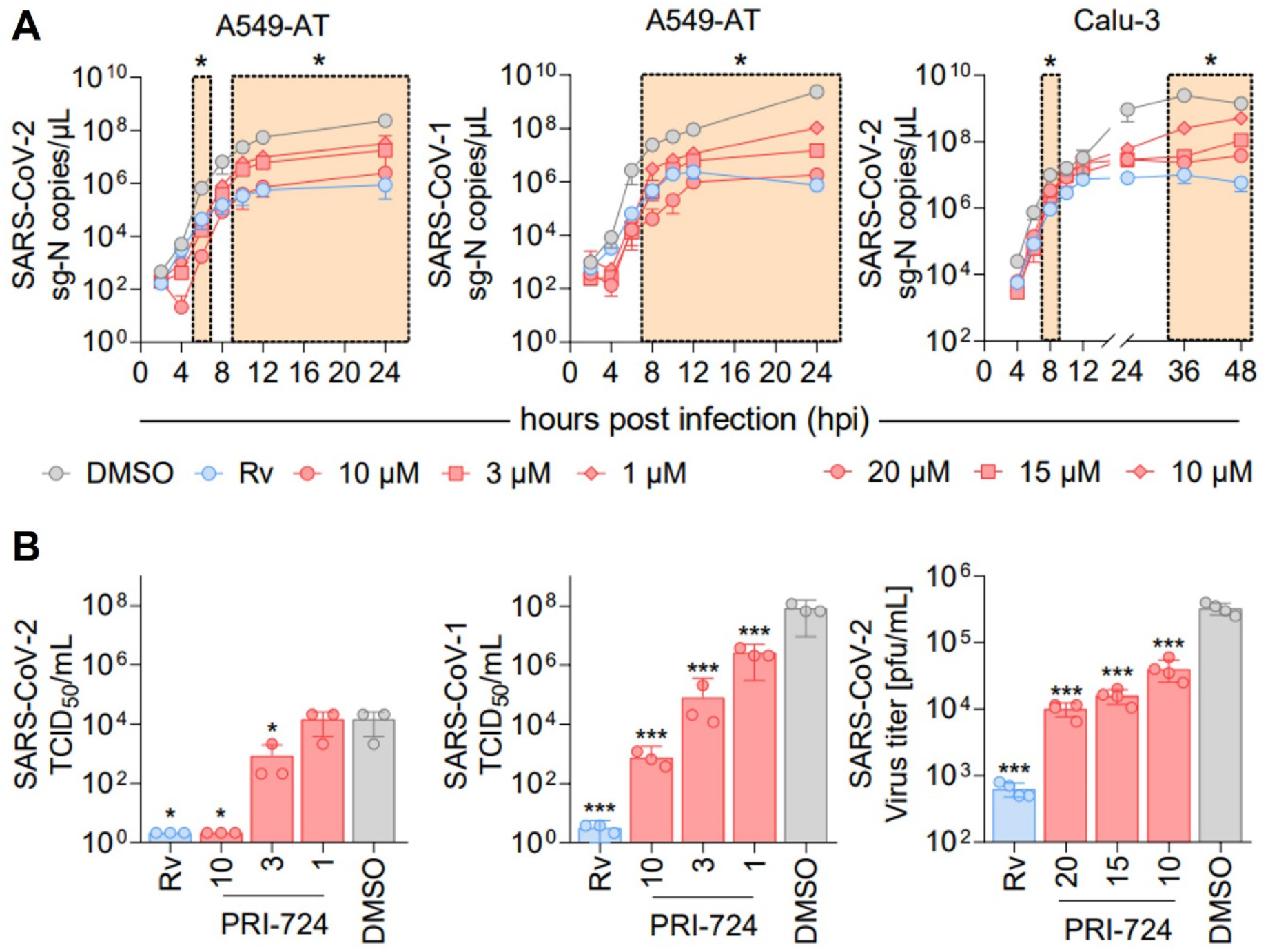
RNA kinetics and quantification of infectious virus upon PRI-724 treatment. A549-AT and Calu-3 cells were treated with 3 μM Remdesivir (Rv, blue) and indicated concentrations of PRI-724 (red; A549-AT: 10 μM (circle), 3 μM (square), 1 μM (rhombus); Calu-3: 20 μM (circle), 15 μM (square), 10 μM (rhombus)), as well as DMSO (grey) and were subsequently infected with SARS-CoV-2 (B.1.617.2 isolate) and SARS-CoV-1 (FFM1 isolate) at MOI=0.1. **(A)** Viral sg-N RNA levels in total RNA lysates over time in A549-AT infected with SARS-CoV-2 (left) and SARS-CoV-1 (middle), as well as in Calu-3 infected with SARS-CoV-2 (right). Data points represent mean and SD of three biological replicates. The experiments were repeated twice with similar results. Significant differences between treatments and control determined by two-way ANOVA are indicated through asterisks; \**p*<0.05 (**Suppl. Tab 8**). **(B)** Infectious SARS-CoV-2 and SARS-CoV-1 virus titers24 hpi in A549-AT determined by TCID_50_ assay (left, middle) and in Calu-3 determined by plaque assay (right). Data points represent mean and SD of three biological replicates. The experiments were repeated twice with comparable results. Significant differences between treatments and control are indicated through asterisks; \**p*<0.05, \*\*\**p*<0.001.

To quantify infectious virus in supernatant, TCID_50_ and plaque assays were implemented (**Fig 5B**). In the supernatant of SARS-CoV-2 infected A549-AT cells, infectious virus was not detectable following treatment with Remdesivir and 10 μM PRI-724. Virus titer was ~17.5-fold reduced upon treatment with 3 μM PRI-724 and no significant reduction was detected with 1 μM PRI-724, which is in line with the obtained RNA levels (**Fig 5A**). SARS-CoV-1 titer was reduced ~10^8^-fold, ~10^5^-fold, 10^3^-fold, and ~31-fold upon treatment with Remdesivir, 10 μM, 3 μM, and 1 μM PRI-724, respectively. Quantification of SARS-CoV-2 in Calu-3 supernatants at 24 hpi by plaque assay revealed 32.5-fold, 20.8-fold, and 8.1-fold reduction in infectious virus titer, when treated with 20 μM, 15 μM, and 10 μM PRI-724, respectively. Overall, a concentration-dependent reduction of SARS-CoV-2 and SARS-CoV-1 vRNA and infectious virus titres upon PRI-724 treatment in A549-AT and Calu-3 cells was observed.

### PRI-724 downregulates β-catenin/CBP target genes *CDC25A, BIRC5* and cyclin D1 protein levels and upregulates β-catenin/p300 target c-Jun

After illuminating the protective *in vitro* properties of PRI-724 during coronavirus infection in A549-AT and Calu-3 cells, we sought to investigate the underlying mechanism. In order to assess β-catenin/CBP activity in A549-AT cells, we implemented a TOPFlash reporter assay (**Fig 6A**), The TOPFlash plasmid contains TCF/LEF binding sites coupled to a firefly luciferase reporter. Nuclear translocated β-catenin binds to TCF/LEF transcription factors to recruit the co-factor CBP to the regulatory element (31) and thereby activates transcription. Treatment with 10 μM, 3 μM, and 1 μM PRI-724 inhibited TOPFlash reporter activity significantly in comparison to the uninduced control. Moreover, induction of TOPFlash activity by 10 mM LiCl was effectively blocked when using the same concentrations. The control construct with mutant TCF/LEF binding sites, showed almost no luciferase activity and no differences among treatments **(Suppl. Fig S3)**.

**Fig 6:**
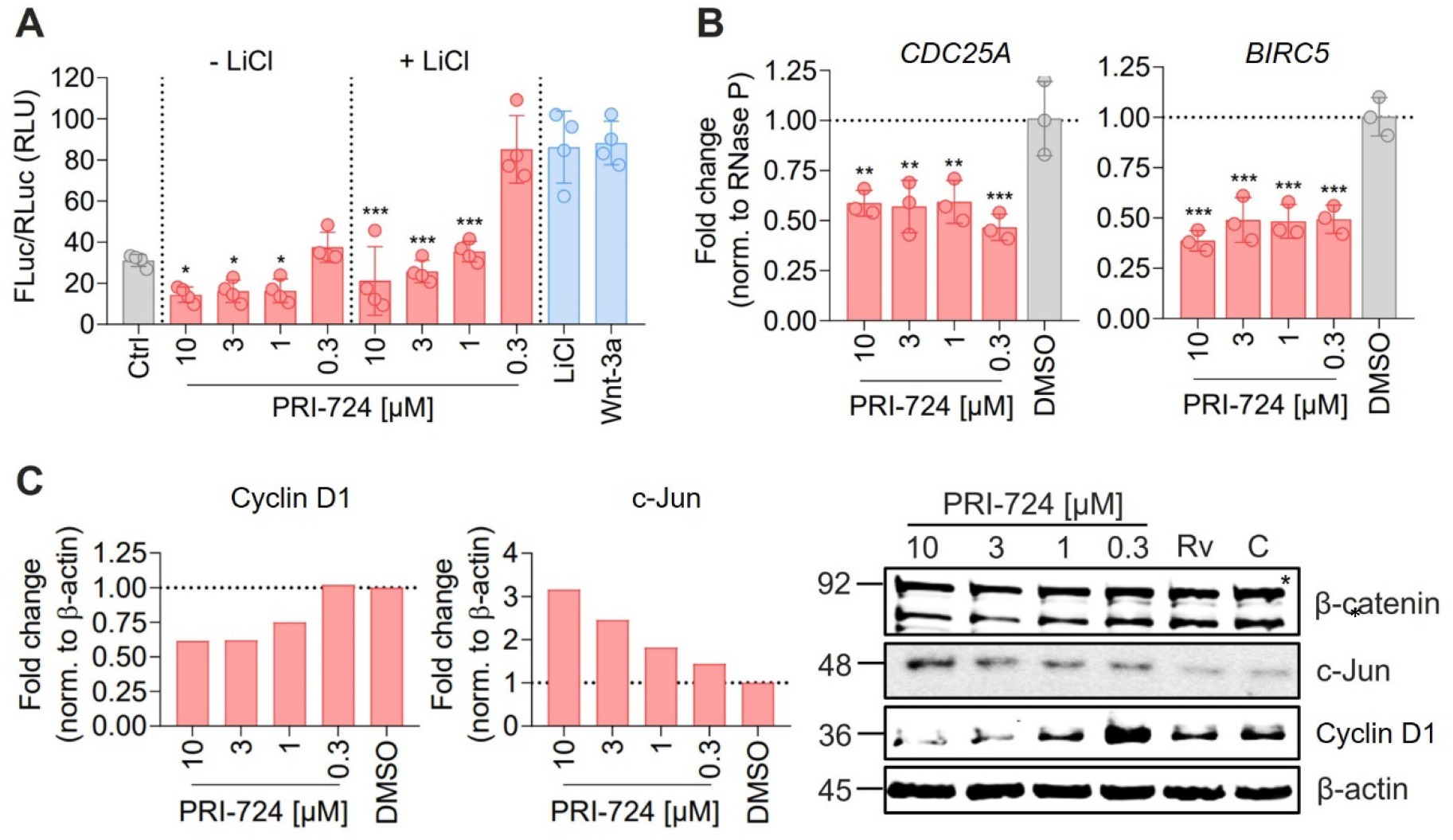
PRI-724 shuffles β-catenin/CBP and β-catenin/p300 activity distribution. **(A)** A549-AT cells were transfected with Super M50 8x TOPFlash and pRL-SV40. After 24 h, cells were treated with 10 mM LiCl, 100 μg/mL rhWnt-3a, and 10 μM, 3 μM, 1 μM and 0.3 μM PRI-724. Luciferase activity was measured after 24 h. Firefly luciferase (FLuc) was normalized to Renilla luciferase (RLuc) for every sample. Data represent mean and SD of three biological replicates. Significant differences are indicated by asterisks; \**p*<0.05, \*\*\**p*<0.001. The experiments were repeated twice with comparable results. **(B)** A549-AT cells were treated with indicated concentrations of PRI-724 and DMSO. 24 h post treatment, RNA was isolated and qRT-PCR was implemented to analyze *CDC25A* (left), *BIRC5* (right), and *RNase P* expression. Data represent mean and SD of three biological replicates. Significant differences are indicated by asterisks; \*\**p*<0.002, \*\*\**p*<0.001. Experiments were repeated twice with comparable results. **(C)** Cells were treated as described in (B). Signals were normalized to β-actin expression. The asterisk designates the predicted band size of β-catenin. Rv – Remdesivir, C – Control.

We further verified the effect of PRI-724 on β-catenin/CBP activity by qRT-PCR and western blot analysis. The expression of the direct target genes *CDC25A* and *BIRC5* were reduced ~2-fold in PRI-724-treated A549-AT cells (**Fig 6B**). Also, t protein levels of cyclin D1 were reduced when treated with 10 μM and 3 μM PRI-724. Reciprocally, β-catenin/p300 target c-Jun was increased in a concentration dependent fashion **(Fig 6C)**. β-catenin expression remained unchanged upon treatments.

### PRI-724 causes cell cycle deregulation in A549-AT cells

It has been reported that PRI-724 causes cell cycle deregulation and block proliferation in several cancer cell lines, such as hepatocellular carcinoma (32), pancreatic (33), and colon cancer (34, 35). We therefore determined whether PRI-724 and/or SARS-CoV-2 infection (MOI 0.01) effects host cell cycle progression. Flow cytometry analysis revealed a significant reduction in G1 (55-60% to 40%) and an increase in S cells (20-22% to 32-35%) upon treatment with PRI-724 compared to DMSO. No significant differences were obtained in G2/M populations. Sub-G1 populations were significantly elevated upon PRI-724 treatment and infected samples showed significantly higher sub-G1 populations than non-infected samples. SARS-CoV-2 infection did not seem to have any significant influence on the cell cycle distribution, although we noticed a slight but significant reduction in the S population of DMSO-treated infected cells compared to non-infected cells. Furthermore, western blot confirmed decreased amounts of survivin and cyclin D1 upon treatment with PRI-724. Also, infected cells treated with DMSO but not with PRI-724 showed slightly increased CDK4 expression (**Fig 7A**). PRI-724-treated A549-AT cells also showed reduced proliferation (**Fig 7B**). To visualize whether PRI-724 induces lateral growth hindrance, a scratch assay was implemented. 10 μM PRI-724 significantly hampered scratch closure at 24 h post treatment. After 24 h, the scratch size remained at ~310 μm over the course of the experiment while the scratch was almost entirely closed up in the control after 72 h (**Fig 7B**).

**Fig 7:**
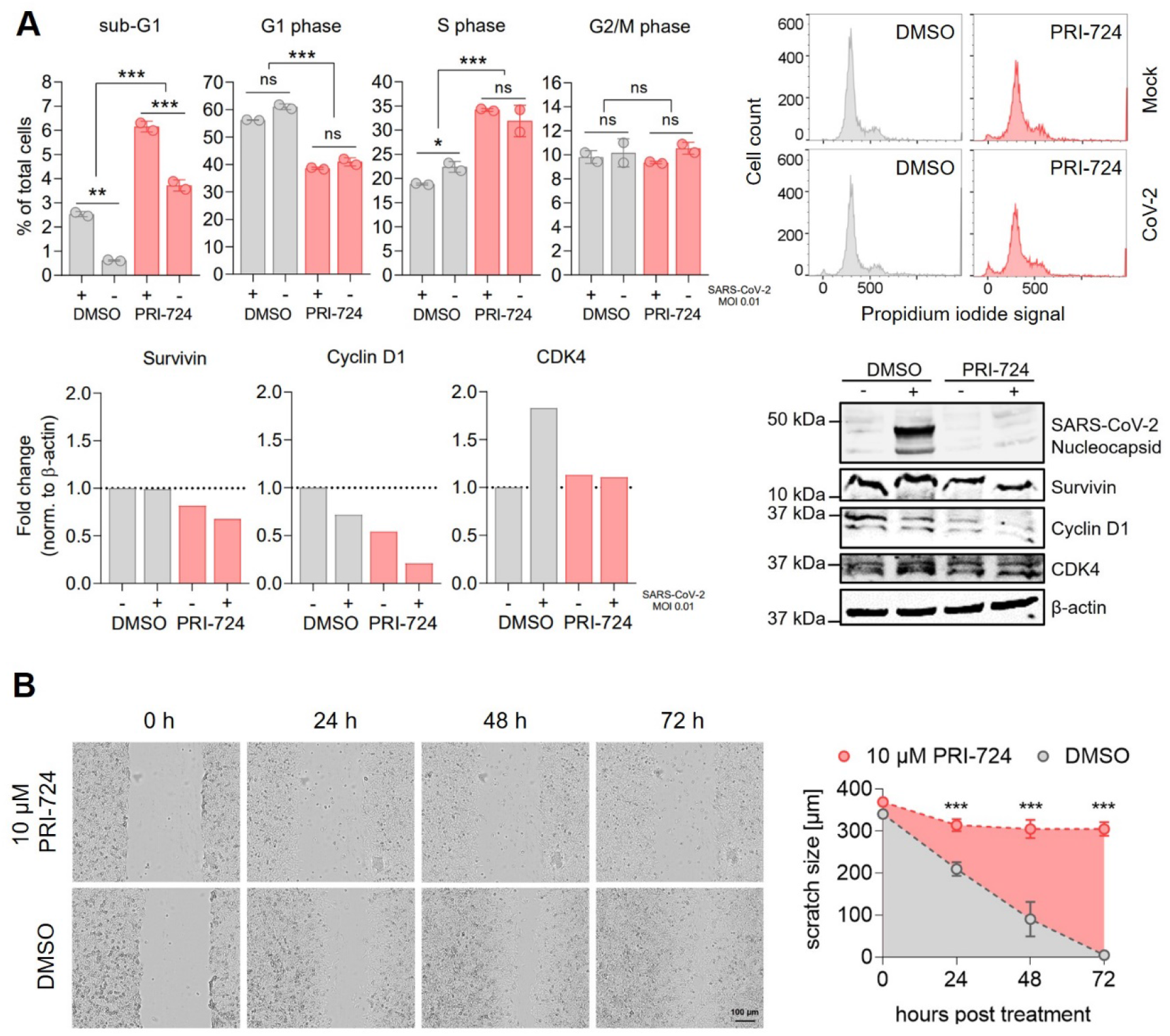
Cell cycle analysis in A549-AT upon treatment with PRI-724. **(A)** A549-AT cells were treated with 10 μM PRI-724 or corresponding amounts of DMSO, and were subsequently infected with SARS-CoV-2 (B.1.617.2 isolate, MOI 0.01). After 24 h, cells were stained with propidium iodide (PI) following flow cytometric analysis. G1, S and G2/M populations were calculated using the Watson model implemented in FlowJo (v10.8.1). Sub-G1 populations were measured manually. Additionally, cell lysates were analyzed by western blot for N protein, survivin, cyclin D1 and CDK4 expression. Expression levels were normalized to β-actin expression. **(B)** Confluent A549-AT cells were scratched prior to treatment with 10 μM PRI-724 or corresponding DMSO amounts. Live cell imaging was performed 0, 24, 48 and 72 hours post treatment to assess scratch healing properties using a Spark^®^ Cyto imaging system. All data represent mean and SD of three biological replicates. Experiments were repeated twice. Significant results obtained by one-way ANOVA are indicated by asterisks; \**p*<0.05, \*\**p*<0.002, \*\*\**p*<0.001.

## Discussion

In this study we provide a comprehensive and robust machine learning-based host factor identification strategy (36) using HDF screens performed for SARS-CoV-2 (16–19) and a drug screen (22). HDF screens of human or African green monkey cells infected with SARS-CoV-2 showed limited overlap in their hits. A better consistency was observed when regarding cellular pathways enriched in observed HDF of the screens. We reason this observation by variances among the different screens due to diverse experimental/technical settings or/and observed biological entities, such as different host cell types or different viral strains. To some extent, such differences may be cleaned out, when regarding a gene set representing a cellular pathway which is essential for the virus. It may contain parts of signaling cascades supporting each other making knocked out genes replaceable in specific conditions. Such interdependencies are highly relevant for discovering drug targets and serves as a promising research venue for future investigations. We took another path here and let machines learn which genes are indispensable, independent of the experimental settings in an automated way. Our machines provided lists of predicted HDF, which reconstruct well the class labels of HDF and non-HDF from unseen data when employing cross-validation. This evidenced that the machines could well capture the data structure in the lists of the gold standards. We followed up on investigating the predictions for their biological content, and particularly their enrichments in known cellular processes and functions. Interestingly, we found a major portion of enriched gene sets being related to development and morphogenesis, followed by gene sets being related to neural processes. This is an intriguing observation in itself, suggesting that particularly cellular processes for development and morphogenesis are hijacked by the virus to facilitate reprogramming of the host cell from cellular proliferation to the proliferation of virus particles.

Focusing on gene sets related to development and morphogenesis, we selected β-catenin as it was highly connected in a constructed protein-protein interaction network, and it was highly expressed during viral infection in our observed cells. β-catenin has been intensively studied, particularly in oncology. It is known for its highly ambivalent roles. In its classical role, it shuttles into the nucleus and acts as a cofactor mainly of p300 or CBP (37). Investigating hematopoietic stem cells (38) and an embryonal murine stem cell line (F9) (39, 40), it was shown that the choice of the interaction partner CBP or p300 matters for the critical decision between proliferation/(pluri)potency and initiation of differentiation. When interrogating publicly available small compound databases, we found PRI-724 and ICG-001 as β-catenin inhibitors. More specifically, they inhibit the interaction of β-catenin with CBP. PRI-724 is an isomer of ICG-001 and the second generation of this drug. We sought to explore the effect of these CBP/β-catenin inhibitors. PRI-724 performed better than ICG-001 **(Fig S1)** and showed an inhibitory effect on pathogenic viruses such as SARS-CoV-2 variants, SARS-CoV-1, MERS-CoV, and IAV, in two different cell culture models. The inhibition doses were cell-type dependent which might be due to varying proliferation rates of the cells and different basal levels of β-catenin and CBP.

Viruses manipulate the cell cycle to generate resources and cellular conditions beneficial for viral replication and assembly. Since PRI-724 is also known to effect cell cycle distribution, we reasoned PRI-724 dependent reductions in virus replication and progeny virus production might be due to cell cycle deregulation. Sui *et al*. observed that SARS-CoV-2 induces dose dependent S and G2/M arrest at the early phase of infection, particularly, when higher MOIs were used, to facilitate virus replication (41), and, in turn, at a higher MOI (0.1), a significant decline of cells in S and G2/M phase at the late phase of infection (24 and 48 hpi). In our experiments we did not observe any significant difference following infection 24 hpi. This is most likely due to lower MOI (0.01) used in our study and is hence in line with the late phase observations from Sui et al. We detected that PRI-724 dependent G1-population decrease and S population increase is independent of the infection and thus it can effectively reduce virus replication and production for different SARS-CoV-2 variants, SARS-CoV-1, MERS-CoV and IAV. Regarding the drug toxicity, the total number of cells was lower after PRI-724 treatment (**Suppl. Fig. S4**) and flow cytometry analysis showed a slight increase in sub G1 population which was significantly pronounced following SARS-CoV-2 infection (**Fig. 7A**). The cell distribution pattern we found differs from the previously published work showing rather an increase in G0/G1 cell population (32, 42) while using ICG-001 and C-82 (active form of PRI-724) when applied to non-infected cells. This observation seems to be cell line or compound type dependent, because another study using colon carcinoma HCT116 cells also found a decrease in G1 and an increase in S cell population following PRI-724 treatment (35).

PRI-724 treatment has been also clinically investigated. PRI-724 was shown to reduce hepatitis C virus-induced liver fibrosis in mice (43). These investigations were followed up in two clinical trials of the same group (phase 1 and 1/2a, respectively) (44, 45). The authors concluded that PRI-724 treatment was well tolerated by patients with HBV and HCV induced liver-cirrhosis and showed improvements of the pathology of concern in several patients. Besides this, blockade of β-catenin/CBP reversed pulmonary fibrosis (46), which is a central COVID-19 complication (47). A recent study confirmed overall higher β-catenin levels in SARS-CoV-2 infected patients and targeting reduced virus shedding *in vitro* (48).

A limitation of our study is that the machine learning procedure bases on screening data of the four initial screens which were at hand when we developed the computational method. New screens have been, and certainly more will be published as means to better understand the mechanisms of this wide spread and evolving virus. As a future outlook, we suggest repeating this computational analysis once, the publication rate of these screens diminished. Furthermore, in this study, our experimental screens based on the observations of two respiratory epithelial cell lines. Further investigations with primary cells are needed to confirm our results.

In conclusion, the machine learning approach brought up interesting connections between viral budding and the reprogramming of host cells in the context of SARS-CoV-2 infection. It may also suit for host factor studies of other viruses and a comparison between different virus entities may lead to intriguing new aspects in virus biology. We observed a strong correlation between the inhibition of CPE, vRNA, and infectious virus production for the tested concentrations in A549-AT and Calu-3 cells, making PRI-724 a promising HDA against SARS-CoV-2 and SARS-CoV-1 in our models. Further investigations, particularly in primary cells is necessary before setting up a clinical trial.

## Materials and Methods

### Gene prioritizations

Two sets of machines were set up to predict HDF, one based on data from CRISPR/Cas9 based knockout screens, and one on data from a drug screen. For defining the gold standard for the first set of machines, the experimental results from four genome-wide CRISPR/Cas9 knockout screens were considered (16–19) (**Tab 2**). The ranking of the screened genes was taken from the supplementary material of the respective studies. The rank product was calculated for each gene for which information was available. The n=500 top ranking genes of this new list were selected and used as the positive class for the classifier, labelled as HDF. The bottom 1,000 genes were used as the negative class (“Non-HDF”). The rest of the genes in the list were used for prediction, i. e. assessing an HDF or non-HDF label. Daniloski *et al*. provided data of two screens with different MOI (0.01 and 0.3). To get a combined list of genes, we calculated the rank product for each gene listed in their score tables and used this combined ranking for our study. In addition, a second gold standard was established based on the screening of 5,632 compounds and their capacity to inhibit the cytopathic effect of SARS-CoV-2 in a Caco-2 cell line (22). In line with the authors of the original study, compounds that blocked viral CPE by 75% were considered as hits. This led to 273 compounds inhibiting SARS-CoV-2 infection without inducing toxicity. Then, their respective targets were identified using the information of different drug databases (Drugbank (49), ChEMBL (50), TTD (51), PharmGKB (52), and BindingDB (53)). By this, 178 gene targets were found and labelled as “HDF”. For the remaining drugs with no inhibitory effect, their 1,881 targets were labelled as “non-HDF”. Effective drugs with more than 11 targets were considered to be “promiscuous”, and their targets were not included in the gold standard.

**Tab 2.**
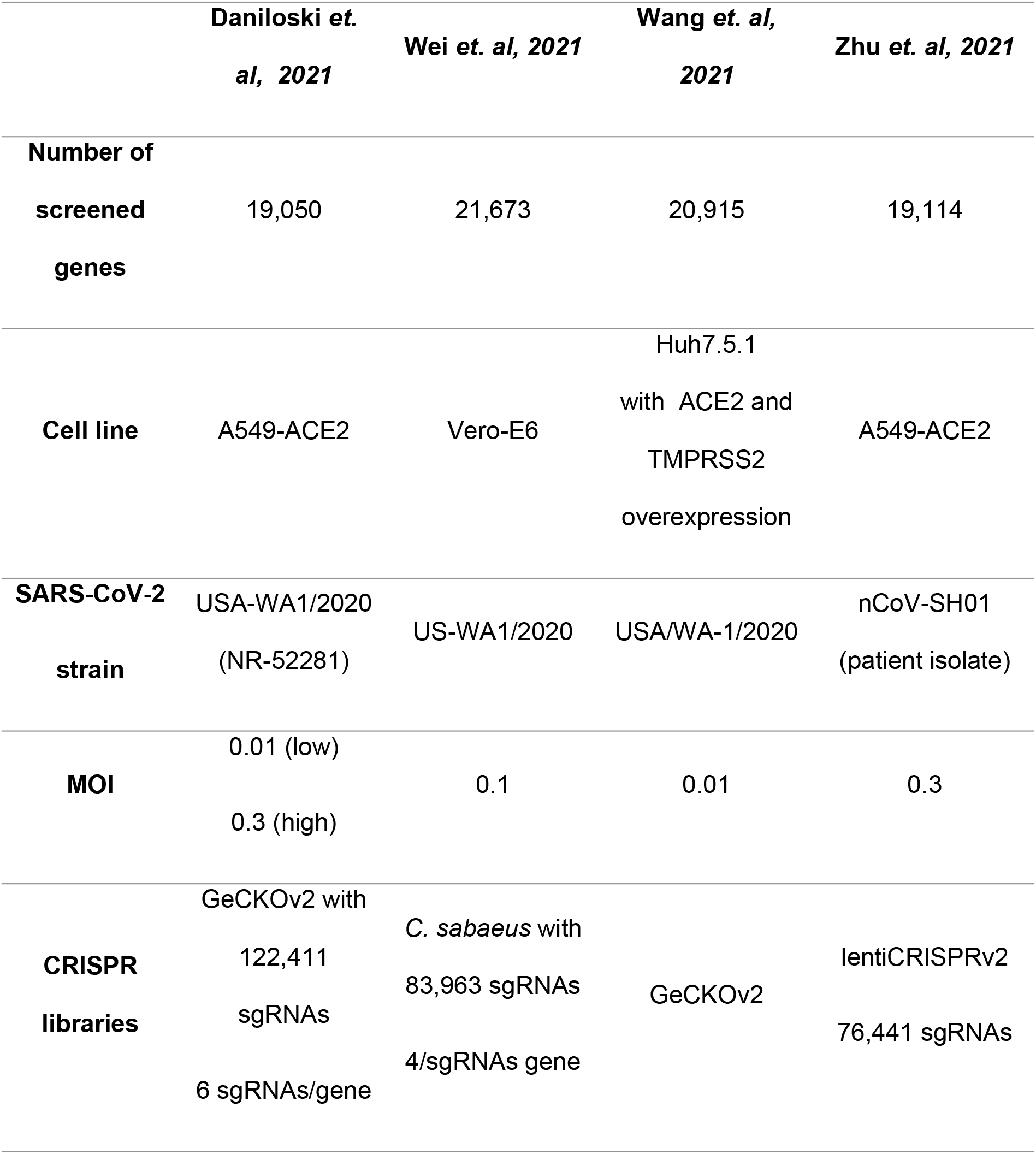
HDF screens serving for a gold standard to train and validate the machines

### Feature generation

The procedure for feature generation was similar as published earlier for essential gene prediction (54, 55, 78). Each gene served as a sample in the machine learning procedure, either labelled as HDF or non-HDF, or was not used for training the classifiers. For each gene, we generated a comprehensive set of 60,593 features comprising the seven categories of (1) gene sequence, (2) protein sequence, (3) protein domains, (4) gene sets from Gene Ontology, (5) conservation, (6) topology in the protein interaction network and (7) subcellular location of the protein. We distinguished between intrinsic and extrinsic features. Intrinsic features denote features, which can be directly derived from gene and protein sequences, whereas extrinsic features describe network topology, homology, cellular localization of the expressed protein and functional (interaction) domains. Employing BioMart (56), we downloaded the gene and protein sequences from Ensembl (v102). Protein and gene sequence features (categories 1 and 2) were calculated with several software tools (seqinR (57), protr (58), CodonW (http://codonw.sourceforge.net), and rDNAse (https://github.com/wind22zhu/rDNAse)) spanning a large range of descriptors as explained in the following. Using seqinR, we calculated the number and percentage of each of the twenty amino acids in a protein. In addition, it was used to calculate other amino acid sequence features including the number of residues, the percentage of each of the physico-chemical classes and the theoretical isoelectric point. Using protr, we calculated the autocorrelation, Conjoint Triad Descriptors (CTD), quasi-sequence order and pseudo amino acid composition. CodonW was used calculating general gene features like gene length and GC content, and frequencies of optimal codons (59) and the number of codons in the coding sequence. rDNAse calculated additional gene features, like auto covariance, pseudo nucleotide composition, and kmer frequencies (n = 2–7). It also calculated the sequence attribute distribution. Amino acids were grouped into three classes (1) polar, (2) neutral or (3) hydrophobic. The second digit in the feature name accounted for the class. More fine grained, seven attributes comprising (1) hydrophobicity, (2) normalized van-der-Waals volume, (3) polarity, (4) polarizability, (5) charge, (6) secondary structure, and (7) solvent accessibility were represented by the first digit in the feature name. The last three digits described the location of the attribute in the sequence, i.e. either at the beginning of the sequence (000), around the 25% quantile of residues (025), 50% (050), 75% (075), or at the end of the sequence (100). For example, seq.attribute.distribution 42100 is the sequence attribute of amino acids being polarizable (4), neutral (1) and are located at the end of the sequence (100). The domain features (category 3) were calculated using BioMart yielding protein family domains (pfam domains), the number of coiled coils structures, prediction of membrane helices, post-translational modifications, β-turns, cofactor binding, acetylation and glycosylation sites, signal peptides and transmembrane helices, and the number and lengths of UTRs. To calculate the gene set features (category 4), gene sets of all terms from Gene Ontology (Biological Process, Molecular Function, Cellular Localization) were obtained from the gene annotation of Ensembl (v102). First, to each gene, its direct neighbors were added using the STRING database (60), and with this list a gene set enrichment test performed employing Fisher’s exact tests. The negative log10 value of the p-value P was used as a feature. Highly redundant (overlapping) gene sets were removed by the following method. Overlap of genes among each pair of gene sets V and W was quantified by Jaccard similarity coefficients,

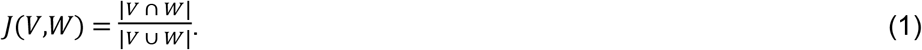

Pairs with J(V,W) above the threshold α=0.3 were included into a model and represented as an undirected graph G(X,E), in which X were the gene sets and pairs above α the edges E. A Mixed Integer Linear Programming problem was set up,

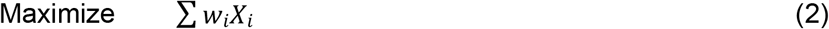

Subject to

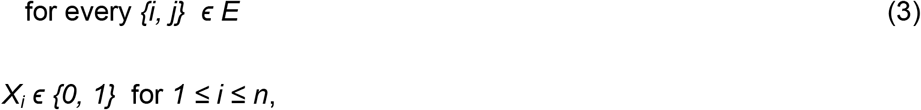

in which w_i_ was the weight of a gene set and calculated from the significance value (P-value P) of the according Fisher’s Exact test,

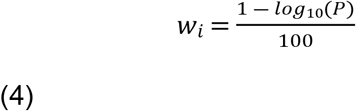

The problem was solved using Gurobi™ (version 7.5.1, www.gurobi.com). By this, up to one gene set of each pair with high overlap was selected and the gene set with the higher enrichment was privileged. Gene conservation (category 5) was assessed by sequence alignment of the protein sequence of each gene with all proteins listed in RefSeq (61) using PSI-Blast (62). The number of obtained proteins with E-value cutoffs from 1e-5 to 1e-100 served as feature values of the respective features. For category 5, typical topology features like degree, degree distribution, several centrality descriptors and Page rank was obtained using protein interaction data (n=697,802 interactions) from the BIOGRID database (77). In addition, we assembled a second network considering also the interactions between SARS-CoV-2 proteins and proteins of the human host cell. For this, in total 22,586 interactions between SARS-CoV-2 and human proteins were obtained from the BioGRID COVID-19 Coronavirus Curation Project (63). Viral proteins were added to the generic network and an interaction was added to the network between a host protein and a viral protein if the link was listed in the protein interactions between SARS-CoV-2 and human proteins. Using both protein interaction networks, in summary, 24 topological features were generated using the igraph R package for network analysis (https://igraph.org). Subcellular localization (category 7) was derived using the prediction software DeepLoc (64), which assigns probability scores to eleven eukaryotic cellular compartments (cytoplasm, nucleus, mitochondria, ER, Golgi apparatus, lysosome, vacuole and peroxisome, plasma membrane, extracellular, chloroplast).

Furthermore, 69 features from experimental data derived from gene expression, proteomics and phospho-proteomics profiles of Caco-2 cells post infection with SARS-CoV-2 at different time points were used. Proteomics data was taken from a previous publication (65) and z-score transformed peptide counts for each time point post infection (0, 2, 6, 10, 24 hrs, MOI=0.01) *versus* time point zero taken. In line, own gene expression data from Caco-2 cells infected with SARS-CoV-2 (FFM1, Wuhan wildtype) (0, 3, 6, 12 and 24 hpi, MOI=0.01). For both datasets, as features, log_10_ differential expression values between each time point post infection and time point zero (no infection) was taken. The phosphoproteomics data was taken from a previous study (SARS-CoV-2 infected Caco-2 cells, 24 hpi, MOI = 1) and features calculated similarly as for the proteomics data (66).

### Machine learning pipeline

Each feature was normalized using z-score transformation. We trained and validated two different sets of machines, one based on the gold standard of the knockout screens (knockout screens based classifier) and one based on the gold standard from the drug screen (drugs screen based classifier). The machine learning procedure was the same for both. For each, a 5-fold cross-validation (CV) was performed in which 4/5 of the data was used to train the model, and the remaining 1/5 of the data (validation set) was used to assess the performance leaving the testing set unseen. In addition, we repeated these cross-validations five times and averaged the results over these five independent runs. The first step in the training procedure was feature selection. For this, Least Absolute Shrinkage and Selection Operator (LASSO) regression was performed using the “glmnet” R package (cv.glmnet function with parameters alpha = 1, type.measure = “auc”). In a second feature selection step, highly correlating features with Pearson’s correlation coefficients r > 0.70 were removed. On average 155 features remained. These features were fed into a Random Forest (RF) classifier using the caret package in R (tuneLength=10, metric = “Kappa”, nthread=20, ntree= 500). All trained models were used to predict the remaining set of 20,007 genes (excluding HDF and non-HDF from the gold standard) to find novel HDF candidates, and to confirm or reject the HDF annotation of the gold standard. We defined the top 10% of the predictions from a voting scheme as predicted HDF (predHDF) of the classifiers, i.e. of the knockout based classifier and of the drug based classifier, leading to 2,182 and 1,989 predHDF, respectively. To evaluate the performance of the learned models, the area under the receiver operating characteristic (ROC-AUC) and area under the precision-recall curve (PR-AUC) of the results from the validation data was calculated using the PRROC package in R. Average and standard variances of the performance estimates were calculated using the results of the individual cross vaildation runs. Most discriminative features of the model were obtained using the varImp function from the caret package which ranked the features based on their impact on the decision trees accuracy.

### Gene set enrichment analyses

The lists of predHDF were investigated for enriched gene sets using the R package gprofiler2 (67). The statistical significance threshold for all GO terms was set to a false discovery rate P < 0.05. To focus on non-too-general and non-too-specific GO terms, only terms with more than 10 and less than 200 genes were considered. Redundancy of the terms was removed as described above (section 4.2) using α=0.4.

### Drug selection

Aiming for repurposing known drugs including clinically approved drugs for the identified targets, a manually assembled data repository of drugs was set up based on seven public drug databases: Drugbank, ChEMBL, TTD, PharmGKB, BindingDB, IUPHAR/BPS and DrugCentral (49–53, 68, 69). The databases provide information about the mechanisms of actions of the drugs, drug-target interactions and therapeutic indications. We considered only drugs approved by either of the following regulatory agencies: Food and Drug Administration (FDA), Health Canada, European Medicines Agency (EMA), and Japan Pharmaceutical and Medical Devices Agency (PMDA), and drugs currently tested in clinical trials.

### Tissue culture and viruses

Lung adenocarcinoma cell line expressing additional ACE2 and TMPRSS2 copies (A549-AT) (70) was cultured in Minimum Essential Medium (MEM) supplemented with 10% (v/v) fetal calf serum (FCS), 100 U/mL penicillin + 100 μg/mL streptomycin (Thermo Fisher; Waltham, Massachusetts, USA) and 2% (v/v) L-glutamine. Calu-3 was cultured in Dulbecco’s Modified Eagles Medium (DMEM)/Ham’s F12 (Thermo Fisher; Waltham, Massachusetts, USA) supplemented with 10% (v/v) FCS, 100 U/mL penicillin + 100 μg/mL streptomycin. Vero E6 was cultivated in DMEM (Thermo Fisher; Waltham, Massachusetts, USA) supplemented with 10% (v/v) FCS and 100 U/mL penicillin + 100 μg/mL streptomycin. Cells were incubated at 37°C, 5% CO_2_. If not otherwise indicated, culture reagents were purchased from Sigma (St. Louis, MO, USA).

Coronaviruses used in this study were isolated from throat swabs obtained from infected individuals and were grown in tissue culture (**Tab 3**). Cell-free aliquots were stored at −80°C and titers were determined by tissue-culture infectious dose (TCID_50_) assay in A549-AT cells.

**Tab 3:** Clinical coronavirus isolates used for infection experiments in this study

| Virus | Specification | GenBank ID |
| --- | --- | --- |
| SARS-CoV-2 | B, FFM5/2020 | MT358641 |
| SARS-CoV-2 | B.1, FFM7/2020 | MT358643 |
| SARS-CoV-2 | B.1.1.7, FFM-UK7931/2021 | MZ427280 |
| SARS-CoV-2 | B.1.351, FFM-ZAF1/2021 | MW822592 |
| SARS-CoV-2 | B.1.617.2, FFM-IND8424/2021 | MZ315141 |
| SARS-CoV-2 | P.2, FFM-BRA1/2021 | MW822593 |
| SARS-CoV-2 | B.1.429, FFM-CALsprt/2021 | MZ317896.2 |
| SARS-CoV-2 | B.1.617.1, FFM-IND5881/2021 | MZ315140 |
| SARS-CoV-2 | B.1.1.529 BA.1, FFM-ZAF0396/2021 | OL800703 |
| SARS-CoV-2 | B.1.1.529 BA.2, FFM-BA.2-3833/2022 | OM617939 |
| SARS-CoV-2 | B.1.1.529 BA.5 | NA |
| SARS-CoV-1 | Frankfurt 1 | AY291315 |
| MERS-CoV | EMC/2012 | JX869059 |
| Influenza A | H5N1/Hongkong/213/03 | NA |

According to the committee for Biological Safety (ZKBS), the entirety of infectious work presented in this study was conducted under BSL-3 conditions. Viruses used in this study and corresponding references are listed in Tab 3.

### Inhibitors and chemicals

Compounds used in this study comprise Remdesivir (#HY-104077), ICG-001 (#HY-14428) (MedChem Express; Monmouth Junction, New Jersey, USA), C-82 (#S0990) and PRI-724 (#S8968) (Selleck Chemicals; Houston, Texas, USA). Compounds were resuspended in DMSO (Carl Roth; Karlsruhe, Germany) and aliquots were stored at −80°C. Recombinant human Wnt-3a (rhWnt-3a) (#5036-WN, R&D Systems; Minneapolis, Minnesota, USA) was resuspended according to manufacturer’s instructions and stored at −80°C.

### Compound treatment and infection assay

Inhibitors were diluted serially on confluent A549-AT or Calu-3 cells in 1% (v/v) serum MEM and were subsequently infected at an MOI of 0.1 with viruses for 48 h. Cells were fixed with 3% (v/v) paraformaldehyde (PFA) for 20 min and stored at 4°C in PBS until analysis. Cytopathic effects were quantified by confluency measurement using a Spark® Cyto 400 mulitmode plate reader (Tecan Group Ltd.; Zürich, Switzerland).

### Scratch assay

Scratch assay was implemented to visualize growth hindrance in A549-AT cells upon treatment with PRI-724. 6 · 10^5^ cells were seeded in a 12-well format one day prior to assay. After applying scratches with a pipette tip, cells were washed with 1x PBS and treated with 10 μM PRI-724 and the respective amount of DMSO in 1% FCS MEM. Cells were subjected to live cell imaging 0, 24, 48 and 72 hours post treatment using a Spark® Cyto 400 mulitmode plate reader (Tecan Group Ltd.; Zürich, Switzerland).

### Live cell imaging

In order to monitor the effectiveness of PRI-724 in terms of blocking syncytia formation and cytopathic effects in general, live cell imaging was carried out in A549-AT cells in a 96-well format. 3.5 · 10^4^ cells were seeded and one day after were treated with 10 μM, 3 μM, 1 μM and 0.3 μM of PRI-724 just prior to infection. DMSO and 3 μM Remdesivir served as negative and positive inhibition controls, respectively. Infection with SARS-CoV-2 B.1.617.2 and SARS-CoV-1 FFM1 was carried out at an MOI of 0.1. Live cell imaging was performed by using a Spark® Cyto 400 mulitmode plate reader (Tecan Group Ltd; Zürich, Switzerland) with hourly measurements of confluence and surface roughness for 48 h.

### Immunofluorescence

A549-AT cells were infected with SARS-CoV-2 B.1.617.2, SARS-CoV-1 Frankfurt-1 and IAV H5N1/Hongkong/213/03 at an MOI of 0.1 for 18 h. Post fixation with 3% PFA, cells were washed and permeablized using ice-cold MeOH. Epitopes were saturated by applying blocking solution (20% (v/v) goat serum, 2% (w/v) BSA, 0.3 M glycine, 0.1% Tween-20, 0.002% thimerosal) for at least 1 h. Primary antibodies (**Suppl. Tab S10**) were diluted 1:1000 in blocking solution containing only 1% (v/v) goat serum and were then incubated on cells O/N at 4°C. Cells were washed three times with 1x PBS and were subsequently incubated with secondary anti-IgG-A488 (1:1000) together with 0.2 μg/mL DAPI and for 1 h. Fluorescence imaging was performed with an Operetta CLS™ High Content Analysis System (Perkin Elmer; Waltham, Massachusetts, USA) using integrated Harmony^®^ software v4.9.

### RNA kinetics and RT-qPCR

For assessing intracellular virus RNA replication upon treatment with PRI-724, medium of infected cells was removed 2 hpi and wells were washed three times with 1x PBS and were supplied with fresh 1% serum MEM. Lysates were collected 2, 4, 6, 8, 10, 12 and 24 hpi in A549-AT and 2, 4, 6, 8, 10, 12, 24, 36 and 48 hpi in Calu-3. Samples were prepared and RNA was extracted using RNeasy QIAcube HT (Qiagen; Hilden, Germany) kit according to manufacturer’s instructions. RNA was stored at −20°C until analysis. qRT-PCR was carried out using Reliance One-Step Supermix (Bio-Rad; Hercules, CA, USA) to measure total viral N RNA and sg-N RNA (**Suppl. Tab S9**, P1-P12) in a multiplex experiment (71, 72). Quantifications of cellular genes *CDC25A, BIRC5, RNase P* (**Suppl. Tab S9**, P13-P19) (73) were carried out using Luna Universal One-Step RT-qPCR Kit (New England Biolabs; Frankfurt (Main), Germany) according to manufacturer’s protocol.

### TCID_50_ assay and plaque assay

For the quantification of infectious virus, supernatants were snap-frozen at −80°C. After thawing, supernatants were diluted on confluent Vero E6 in 96-well plates. Four days post infection, dilutions were analysed for CPE development. TCID_50_ was calculated according to the Spearman & Kärber algorithm. For plaque assays, supernatants were diluted on confluent Vero E6 in 24-well plates. After 1 h, supernatants were coated with overlay medium (1.5% methyl cellulose (w/v) (Merck KGaA; Darmstadt, Germany), 1x MEM, 1% (v/v) FCS, 100 U/mL penicillin + 100 μg/mL streptomycin). When plaques reached sufficient size, cells were washed and fixed with 3% PFA for 20 min. Cells were stained with CV staining solution (1% (w/v) crystal violet, 20% (v/v) MeOH).

### Luciferase assay

A549-AT cells were seeded in a 96-well format and were transfected with 0.1 ng of pRL-SV40 (Addgene; #27163) together with either 0.1 μg M50 Super 8x TOPFlash (Addgene; #12456) or M51 Super 8x FOPFlash (TOPFlash mutant) (Addgene; #12457) using *Trans*IT^®^-LT1 Transfection Reagent (Mirus Bio LLC; Madison, Wisconsin, USA). After 24 h, cells were treated with either of the following substances: 10 mM LiCl (Merck KGaA; Darmstadt, Germany), 100 μg/mL rhWnt-3a, 10 / 3 / 1 / 0.3 μM PRI-724. Additionally, cells were treated simultaneously with LiCl and PRI-724. After 24 h, cells were analysed for luciferase activity using a Dual Luciferase Reporter Assay Kit (Promega; Madison, Wisconsin, USA) according to manufacturer’s instructions.

### Western blot

For preparation of protein extracts, cells were lysed using RIPA buffer (150 mM NaCl, 50 mM Tris/HCl pH 8.0, 1% (v/v) Triton X-100, 0.5% (v/v) sodium deoxycholate, 0.3% (v/v) SDS, 2 mM MgCl_2_). Buffer was supplemented with 5 mM NaF, 1 mM Na_3_VO_4_, 20 mM β-glycerophosphate, 1x cOmplete Mini, EDTA-free protease inhibitor cocktail (Merck KGaA; Darmstadt, Germany) and 1 U/mL Pierce™ Universal Nuclease (Thermo Fisher; Waltham, Massachusetts, USA). Lysates were stored at −20°C. Protein concentrations were determined using DC Protein Assay Kit (Bio-Rad; Hercules, CA, USA). Proteins were separated by SDS-PAGE and were blotted onto an Amersham™ Protran^®^ nitrocellulose membrane (Merck KGaA; Darmstadt, Germany) for 1.5 h using const. 120 V. Blots were washed and blocked for at least 1 h using 5% (w/v) BSA/TBS-t (150 mM NaCl, 20 mM Tris, 0.1% (v/v) Tween20). Primary antibody (**Suppl. Tab S10)** incubation was performed overnight at 4°C in 5% BSA-TBS-t. After washing, secondary antibody was incubated for 1 h in 5% BSA-TBS-t. Blots were imaged using CLx imaging device (LI-COR; Lincoln, Nebraska, USA).

### Cytotoxicity measurement by fluorescence microscopy

Confluent A549-AT cells were treated with 10 μM, 3 μM, 1 μM, 0.3 μM PRI-724 and with the highest corresponding amount of DMSO for 48 h in 1% MEM. Cells were fixed with 3% PFA and stained with 0.5 μg/mL Hoechst 33342. Imaging was performed with Olympus IX 81 scanning a total of 9 fields/well with a 3×3 scatter at a total magnification of 10x (pixel size = 0.645 μm). Screening was performed using ScanR Acquisition v2.4.0.13 software.

The analysis was performed using KNIME and KNIP (74). The images were filtered using a gaussian convolution (σ = 2 px). A consecutive image thresholding assigned pixels with values greater than the image-mean to the foreground. Cells were detected and labeled by connected component analysis. Cell clumps were afterwards split into single cells using the Wählby algorithm (75). The resulting components were filtered based on their size, components too large or too small were rejected as they most likely do not depict cells. Cell count was measured for the different PRI-724 concentrations (**Suppl. Fig S4)**.

### PI staining and flow cytometry

In a 6-well plate, confluent A549-AT cells were treated with 10 μM PRI-724 and the corresponding amount of DMSO for 24 h and were infected with SARS-CoV-2 (B.1.617.2 isolate, MOI 0.01) in 1% MEM. After that, cells were gently detached and fixed in ice-cold 70% ethanol for 1 h. Cells were washed three times with FACS-PBS (2% (v/v) FCS, 0.1% NaN_3_) and were then stained overnight at 4°C by applying PI staining buffer (50 μg/mL PI, 2% (v/v) FCS, 200 μg/mL Monarch^®^ RNase A (New England Biolabs; Frankfurt (Main), Germany), 0.1% Igepal CA-630). Cells were analysed for PI staining using FACSVerse™ Flow Cytometer (BD Biosciences; Mississauga, ON, Canada).

### Statistics for experiments

The curve fittings presented for dose-responses to Remdesivir and PRI-724 presented in Fig 3A and in suppl. Fig S1 were performed by applying robust non-linear regression comprising a sigmoidal 4-parameter model for a total of three biological replicates per concentration:

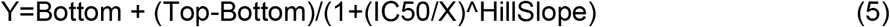

The exact IC_50_ values calculated from these analyses are listed in Fig 3A and in suppl. Tab S6/S7). For simple group comparisons such as data presented in Fig 3B, Fig 4, Fig 5B and 5C, Fig 6A and 6B as well as Fig 7, one-way ANOVA was generally applied to identify significant differences according to treatments. In all tests, equal variances according to the mean were assumed and further analyses were not corrected for multiple testing (Fisher’s LSD test). Results were rated significant when *p*<0.05. For RNA kinetics depicted in Fig 5A, two-way ANOVA was applied to identify significant differences in vRNA levels according to the timepoint of collection and the sort of treatment. Again, equal variances according to the mean were assumed and further analyses were not corrected for multiple testing (Fisher’s LSD test). Results were rated significant when *p*<0.05.

## Acknowledgements

MAK and AH were supported by Research Grants of Merck KGaA, Darmstadt, Germany, TT is funded by Willy Robert Pitzer Foundation. MAK and TT were supported by the Federal Ministry of Education and Research (BMBF), Germany, FKZ: 01KI2015OB, MO by the Federal Ministry of Education and Research (BMBF), Germany, FKZ: 01EO1502 (CSCC), 13N15711 (LPI) and 01KI2015OA (SARSiRNA) and TB by the Deutsche Forschungsgemeinschaft (KO-3678/5-1).

## Authors‘ contributions

TT, HE and RK designed the concept for the study. AVG, AH, TB, MO and RK performed the data analysis and machine learning. MK and TT designed and performed the experiments. NB and PW performed the experiments on Cytotoxicity measurement by fluorescence microscopy. All authors revised and approved the manuscript.

